# Deep learning for microscopy without machine learning expertise

**DOI:** 10.64898/2026.04.03.716415

**Authors:** Xinyu Zhou, Shaopeng Wang

## Abstract

Deep learning can extract quantitative measurements from microscopy images that are inaccessible to classical analysis, but developing these models requires machine learning expertise that most imaging scientists do not have. Here we present a framework in which a researcher describes their microscopy problem to a large language model (LLM) agent in under ten minutes of conversation—specifying what they image, what they want to measure, and what success looks like—and the agent autonomously handles the rest: designing physics-based training data, implementing a neural network, training, diagnosing failures, and iterating without human intervention. A researcher can start the agent before leaving the lab; overnight, it tests tens to a hundred model variations, each one an experiment that would otherwise demand active attention. We validate the framework across six microscopy modalities and four problem types. On the BBBC039 nuclear segmentation benchmark, the agent autonomously trains a U-Net with 3-class semantic segmentation and morphological post-processing, achieving pixel-level Dice of 0.97 and object-level F1 of 0.84—within 7% of the published baseline—while diagnosing a data pipeline bug that no amount of hyperparameter tuning could resolve. On single-protein holographic microscopy, the agent reads a published paper, designs a simulator, and develops an optimized model in a single session. On PatchCamelyon histopathology classification, the agent autonomously evolves through four optimization phases—from scratch training through transfer learning and regularization to inference-time ensembling—completing 97 iterations on 262,144 images to reach 89.3% test accuracy and 96.3% AUC, nearly matching the published rotation-equivariant baseline. This framework enables microscopy researchers to use deep learning-based image analysis without machine learning domain knowledge.

## Introduction

Deep learning methods for microscopy have matured rapidly. Neural networks now match or exceed classical algorithms for single-molecule localization[1, 2], particle tracking[3], diffusion classification[4], and super-resolution reconstruction. Yet adopting these methods remains difficult for the researchers who need them most. A biologist who images cell nuclei, a biophysicist measuring single-protein binding events, or a materials scientist tracking nanoparticle diffusion—each may recognize that a published deep learning approach could transform their measurement, but translating that recognition into a working model requires navigating an unfamiliar stack of software engineering decisions: choosing a framework, implementing an architecture, configuring a training loop, diagnosing CUDA errors, tuning hyperparameters, and validating results. This engineering effort is orthogonal to the domain expertise these scientists already possess, and it commonly takes weeks of trial and error—or is abandoned entirely.

Existing tools address fragments of this problem. Automated machine learning (AutoML) and neural architecture search (NAS) optimize hyperparameters or architecture choices, but require the user to first supply a working data pipeline, training loop, and search space[5]—precisely the infrastructure that non-specialists struggle to build. LLM-based coding assistants can generate code snippets and answer technical questions[6], but they operate reactively, one prompt at a time, leaving the researcher responsible for orchestrating the overall development process. Self-driving laboratory platforms[7] automate experimental parameter optimization but do not extend to developing the computational models that analyze the resulting data. None of these approaches take a domain scientist from “I have microscopy images and a scientific question” to “I have a trained, validated deep learning model” without substantial machine learning engineering in between.

Here we present a framework that closes this gap. A researcher describes their microscopy problem to an LLM agent in a brief interactive session—typically under ten minutes—specifying what they image, what they want to measure, and what success looks like. From this conversation, the agent generates a complete pipeline configuration: physics-based simulation, model architecture, training protocol, and success metrics. The agent then enters an autonomous experimentation loop (the “autoloop”), in which it iteratively implements changes, trains the model, evaluates results against defined metrics, and keeps or reverts each change—cycling without human intervention. When training requires minutes per iteration, an overnight run can yield up to a hundred experiments; when models are larger or data more complex, the same period yields tens of iterations, each one an experiment that would otherwise require a researcher’s active attention.

We validate the framework across six microscopy modalities, five levels of simulation complexity, and four machine learning problem types (Fig. 1) As end-to-end demonstrations, we apply it to three problems: single-protein holographic microscopy, where the agent reads a published paper, designs a physics-informed simulator, and develops a complex-field contrast estimator through 14 autoloop iterations in a single session (Fig. 2); the BBBC039 nuclear segmentation benchmark, where the agent autonomously reproduces published results on real experimental data and, in the process, diagnoses a data pipeline bug that no amount of hyperparameter tuning could have overcome (Fig. 3); and PatchCamelyon histopathology classification, where the agent completes 97 autonomous iterations on 262,144 images, evolving through four optimization phases to nearly match the published baseline (Fig. 4).

**Figure 1:**
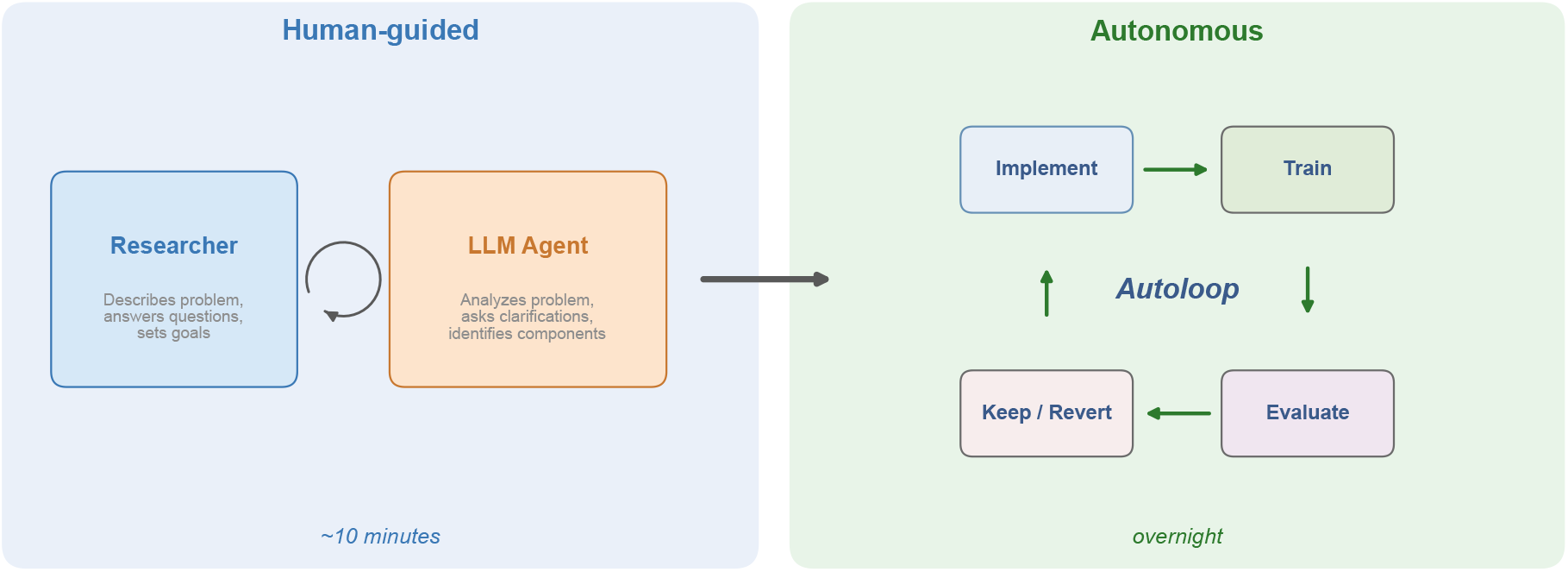
End-to-end workflow for LLM-autonomous ML development. A researcher interactively defines the problem with the LLM agent (left, human-guided): through iterative conversation, they identify the simulation physics, data pipeline, model architecture, success metrics, and experiment plan, producing a structured pipeline configuration. The agent then enters the autoloop (right, autonomous): a bounded cycle of implement–train–evaluate–keep/revert iterations that runs without human intervention. See Extended Data Figs. 1–2 for framework architecture and autoloop cycle details.

**Figure 2:**
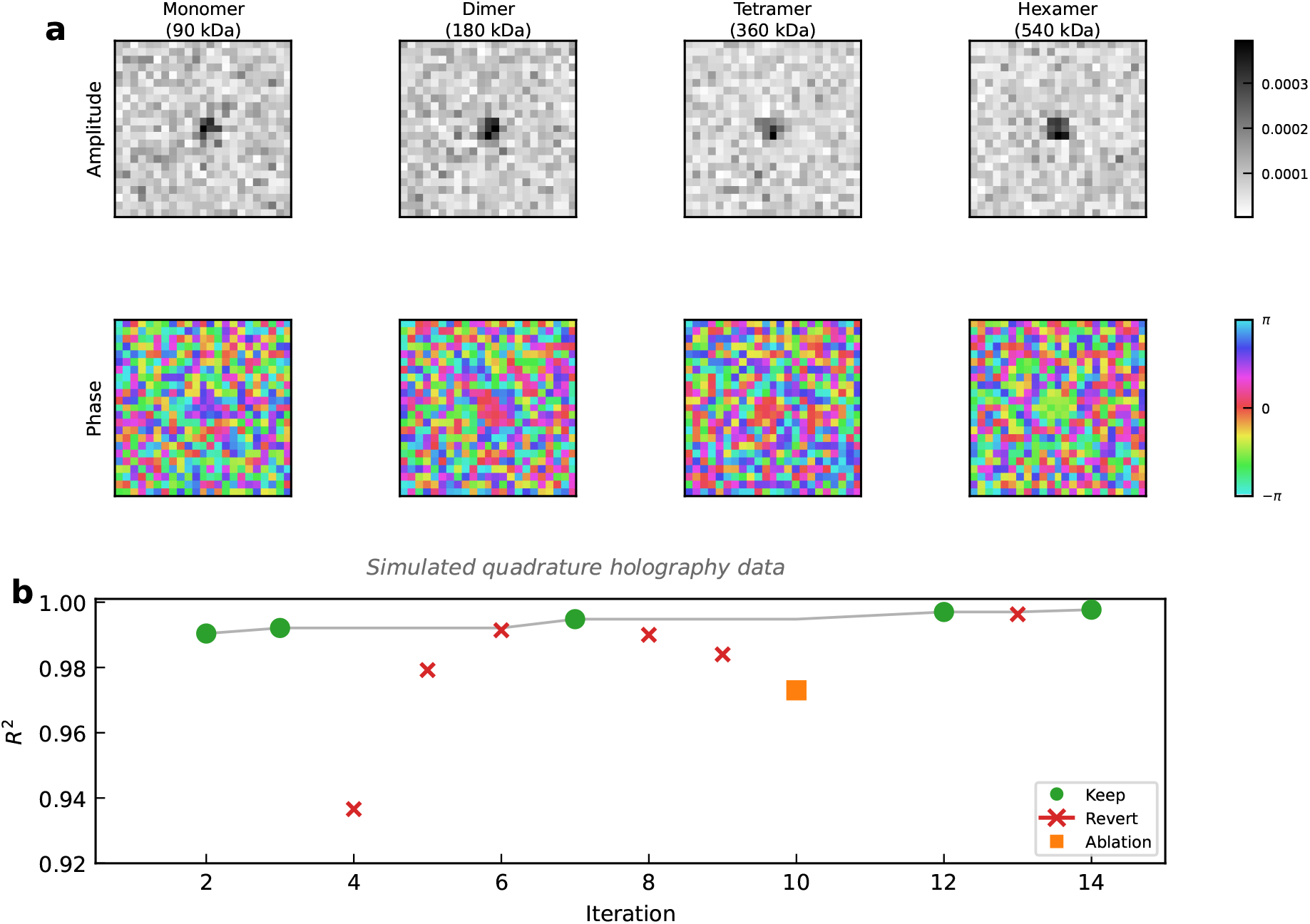
Autonomous development of a complex-field contrast estimator on simulated quadrature holography data. **(a)** Example simulated patches at increasing scattering contrast (monomer to hexamer), showing the amplitude (top, grayscale) and phase (bottom, cyclic colormap) of the reconstructed complex scattered field. Patches are generated using a Jinc PSF model with realistic noise calibrated to the experimental conditions of Thiele et al. **(b)** Autonomous optimization trajectory across 14 agent iterations, showing *R*^2^ on a held-out simulated test set. Kept changes (green circles) and reverted changes (red crosses) are annotated; the single ablation experiment (orange square) tested amplitude-only input.

**Figure 3:**
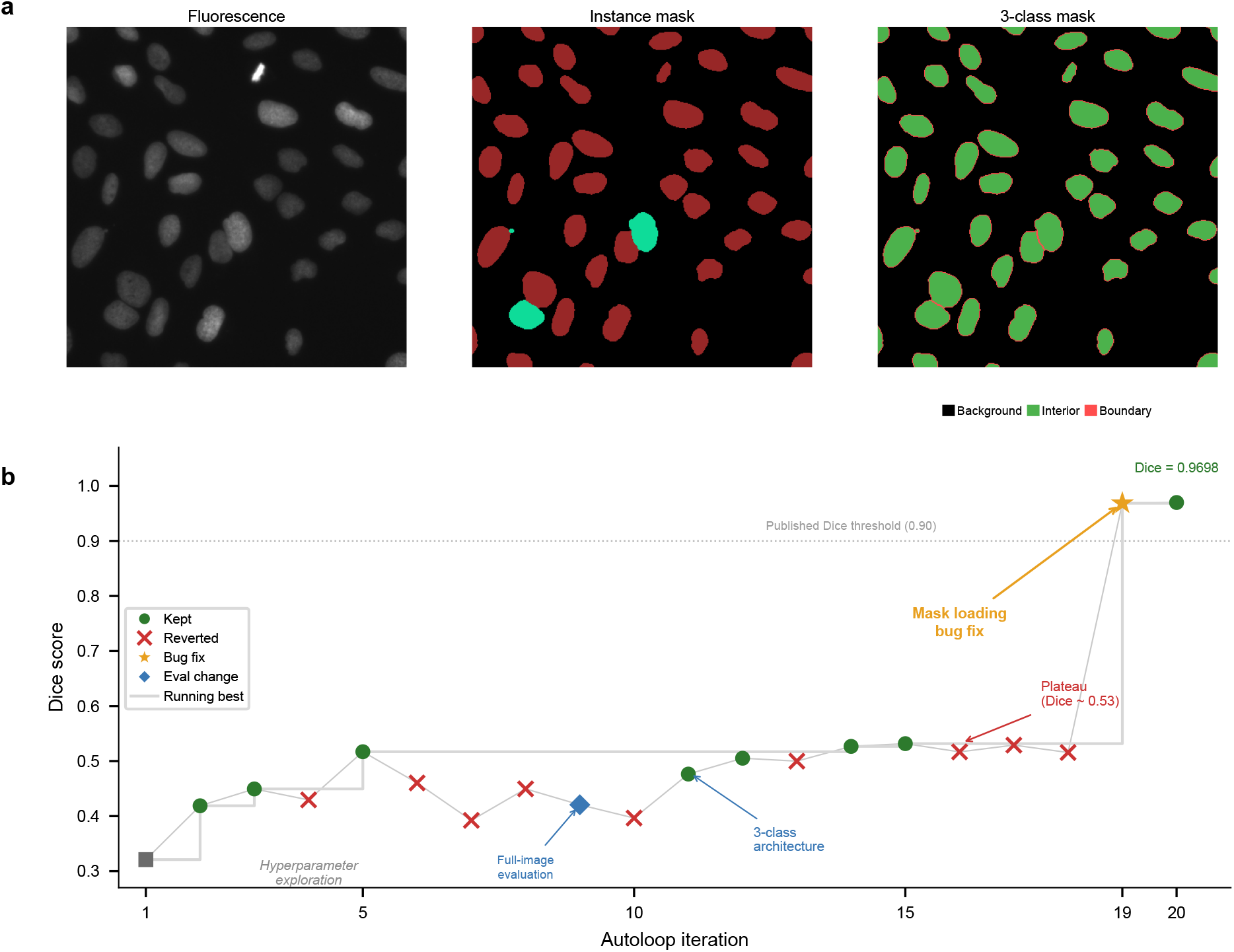
Autonomous benchmark reproduction on the BBBC039 nuclear segmentation dataset. **(a)** Example validation image: raw DAPI fluorescence (left), instance segmentation mask with per-nucleus colors (center), and 3-class semantic mask used for training—interior (green) and boundary (red) classes, following Caicedo et al. **(b)** Autoloop optimization trajectory across 20 iterations. Green circles: kept changes; red crosses: reverted changes; orange star: data pipeline bug fix; blue diamond: evaluation methodology change. The agent autonomously diagnosed a mask loading bug at iteration 19, resolving a persistent plateau at Dice ≈ 0.53 and achieving pixel-level Dice = 0.97 and object-level F1 = 0.84, within 7% of the published baseline. Gray line traces the running best Dice score.

**Figure 4:**
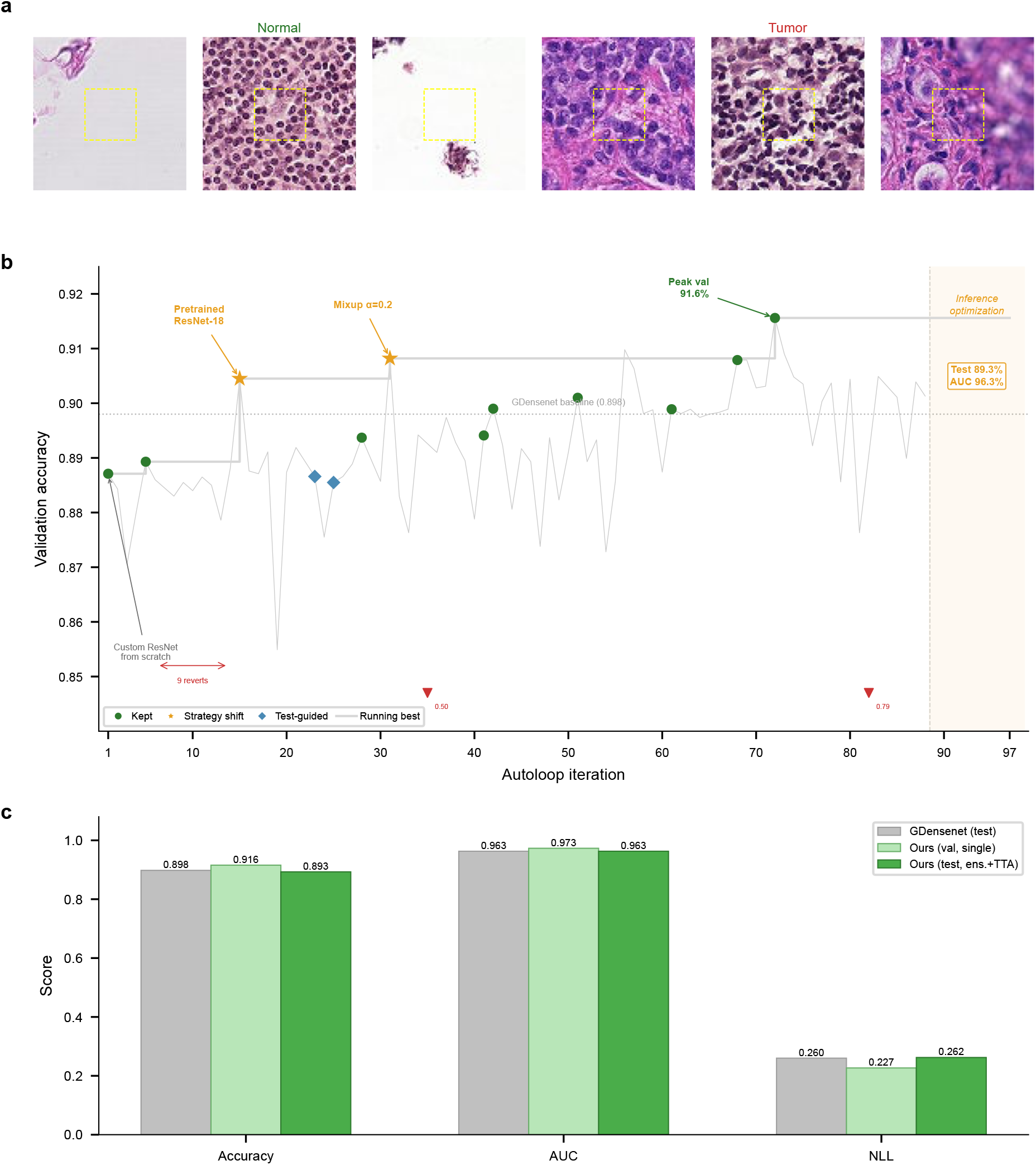
Autonomous benchmark validation on PatchCamelyon histopathology classification. **(a)** Example 96×96 H&E-stained patches: normal tissue (left three) and metastatic tumor (right three). Dashed yellow boxes indicate the center 32×32 region that determines the label. **(b)** Autoloop optimization trajectory across 97 iterations. Green circles: kept changes; orange stars: strategy shifts (pretrained ResNet-18 at iteration 15, mixup at iteration 31); blue diamonds: test-guided decisions. Gray line traces the running best validation accuracy. Four phases are visible: custom ResNet (1–14), regularization and infrastructure (15–51), exhaustive optimization (52–88), and inference-time strategies (89–97). **(c)** Metric comparison: published GDensenet test results[16] (gray), our best single-model validation (light green, iteration 72), and our best test result using top-2 seed ensemble with 8× TTA (dark green, iteration 93). Validation metrics exceed published test values for all three metrics; test AUC matches exactly (96.3%) and test accuracy reaches 89.3%, within 0.5 percentage points of published. For NLL, lower is better.

## Results

### From problem description to trained model

The framework is designed so that a researcher’s only responsibility is describing their problem. In a brief interactive session—typically under ten minutes—the researcher tells the LLM agent what they image, what they want to measure, and what a successful model looks like. From this conversation, the agent generates a complete pipeline configuration: a YAML (YAML Ain’t Markup Language) file specifying the project description, data format, model architecture, success metrics with quantitative thresholds, an experiment plan, and hyperparameter bounds. No code, no environment setup, no architecture decisions.

From this single configuration file, the framework assembles everything the agent needs to work autonomously: a structured protocol defining session startup, task selection, and decision tables; a prioritized experiment plan; and quality gates that prevent broken code from consuming compute. At each session, the agent follows a deterministic routine— checking for pending decisions, reviewing prior results, selecting the highest-priority eligible experiment—and then implements, trains, evaluates, and commits changes without further input. Training metrics are parsed into structured JSON and analyzed against predefined thresholds, producing a verdict for each experiment: SUCCESS, PARTIAL, TIMEOUT, FAILED, or AMBIGUOUS (Extended Data Fig. 1b).

When things go wrong—and they inevitably do—the agent handles errors through a graduated response protocol (Extended Data Fig. 1c). Common failures are self-resolved: the agent halves the batch size upon out-of-memory errors, reduces the learning rate by 10× when training diverges, or extends the time budget after timeouts. When the core experiment remains evaluable despite an issue, the agent logs a caveat and continues. Only when the situation is genuinely ambiguous—when proceeding would risk wasting compute or producing misleading results—does the agent escalate by creating a structured decision document and halting for human review. All autonomous actions are recorded in a triple audit trail: git commits with structured prefixes, a progress log with autonomy tags, and per-experiment logs (Extended Data Fig. 1a).

Across three test problems (gold nanoparticle sizing, nuclear segmentation, quantum dot classification), automated onboarding achieved 100% correctness in generating valid configurations, compared to 87% when the agent operated without the structured setup protocol (Table S1).

### The autoloop: bounded autonomous experimentation

Once configured, the agent enters the autoloop—an autonomous experimentation cycle that runs indefinitely without human intervention (Extended Data Fig. 2). The researcher starts the loop and walks away; results are waiting when they return. Each cycle follows a fixed sequence:

1. **Read** the current experiment log, best metrics, and active strategy.
2. **Decide** the next modification based on a prioritized list of ideas, ordered by expected impact.
3. **Implement** the change in a single, focused code edit.
4. **Gate**: run quality checks (syntax, imports, single-batch forward pass) before committing compute.
5. **Commit** the change to git, creating a revert point.
6. **Train** with a fixed time budget, ensuring each cycle completes within a bounded period.
7. **Extract** metrics from training output by parsing structured log lines.
8. **Evaluate**: compare metrics against the current best. If improved, keep the change and update the best; if not, revert via git.
9. **Log** the iteration result, then return to step 1.

Each iteration takes minutes to over an hour, depending on model size and dataset volume. An overnight run therefore yields tens to a hundred iterations—each one an experiment that would otherwise require a researcher’s active attention to implement, monitor, and evaluate.

Three mechanisms prevent the autoloop from stalling. First, **plateau detection**: after five consecutive reverted iterations, the agent triggers a diagnostic protocol that analyzes failure patterns, checks for data insufficiency, and shifts strategy. Second, **automatic data scaling**: if diagnostics identify data insufficiency as the root cause, the agent doubles the training dataset (up to 4× the original size) and retries. Third, **context management**: the agent monitors its own context window usage and, at a set threshold, saves all state to external files, compresses context, and resumes seamlessly. Since all loop state is persisted in files (experiment log, configuration, git history), context restarts lose no information.

### Generalizability across modalities, complexity levels, and problem types

A central challenge in applying deep learning to microscopy is the scarcity of labeled training data. Our framework addresses this by having the LLM agent autonomously design and implement physics-based simulations. To test generality, we evaluated the agent across three axes.

#### Simulation physics

The agent adapted a common baseline simulation (Brownian dynamics with Gaussian PSF) to six microscopy modalities: TIRF fluorescence, widefield fluorescence, dark-field scattering, interferometric scattering (iSCAT), digital holographic microscopy, and correlative dual-channel imaging. Each adaptation required modality-specific physics—evanescent wave decay, Born–Volf defocus, Mie theory cross-sections, coherent field addition, spherical wave propagation, or multi-population co-localization. No modality-specific code was pre-built; the physics intelligence comes from the LLM’s domain knowledge. The agent achieved 34/34 physics assertions across all six modalities, verified by independent grading agents (Table S2; Supplementary Section S2).

#### Simulation complexity triage

Not all problems can be simulated with numpy. Ve presented five scenarios spanning simple Brownian motion to nucleotide-resolution DNA mechanics. The agent correctly determined when numpy suffices, when external MD engines (HOOMD-blue[8], oxDNA[9]) are needed, and when analytical models are more appropriate—passing 26/26 triage assertions (Table S3; Supplementary Section S3).

#### Problem types

We tested onboarding across four ML problem types: segmentation (U-Net[10], nuclear segmentation), detection (CNN with focal loss[11], nanoparticle detection), classification (ResNet[12], quantum dot counting), and regression (MSE loss, nanoparticle sizing). Each model template is a self-contained single Python file. The setup skill achieved 100% correctness (23/23 assertions) compared to 87% for unguided operation (Table S1; Supplementary Section S5).

### Autonomous model development for single-protein holographic microscopy

To demonstrate the autoloop on a real scientific problem, we applied the framework to single-protein optical holography data from Thiele et al.[13]. The agent was given the published paper and dataset. Without additional guidance, it analyzed the processing pipeline to identify an improvement opportunity, designed a physics-informed simulator, developed a CNN-based estimator, and optimized it through 14 autoloop iterations—all in a single session.

#### Background

Thiele et al. demonstrated single-protein detection using a quadrature holographic microscope with four cameras recording interferograms at phase shifts of 0, *π/*2, *π*, and 3*π/*2. The four images are combined to reconstruct the complex scattered field of individual protein landing events. The published pipeline estimates protein mass from the amplitude only, fitting a parametric Jinc PSF to each landing event. The reported mass resolution is 28 kDa for DynDeltaPRD (90 kDa monomer). The agent identified that the paper measures the full complex scattered field but discards phase information at the estimation stage—a clear opportunity for improvement.

#### Simulation

The agent designed a physics-based simulator modeling the quadrature reconstruction output. Each simulated patch represents the complex scattered field of a point scatterer (Fig. 2a): *E*_scat_(*x, y*) = *c* · *e*^*iϕ*^ · PSF(*x, y*) + noise, where *c* is the scattering contrast and PSF is a Jinc function with *σ* = 2.13 pixels matching the diffraction limit at 450nm, 1.1 NA, and 117.2nm pixel size. The noise model combines a fixed floor (6 × 10^−6^) with a signal-dependent component, matching the post-averaging shot noise. 10,000 training and 2,000 validation patches were generated with contrasts sampled log-uniformly over [2×10^−5^, 5×10^−4^], spanning the full protein mass range.

#### Baseline reproduction

Before developing improvements, the agent reproduced published baseline numbers from the experimental AuNP calibration data (20, 40, 60 nm gold nanoparticles with Mie theory cross-sections), confirming: contrast-to-cross-section conversion factor 28.7 nm (published: 28.1 ± 0.4 nm), contrast-to-mass slope 6.05×10^−7^ per kDa (published: ∼6.0×10^−7^), and specific polarizability 217 Å^3^/kDa (published: 239 ± 4 Å^3^/kDa).

#### Model

The agent developed a CNN taking 2-channel input (real and imaginary parts of the reconstructed scattered field, 23×23 pixels) and predicting scalar log-contrast. Two physics-informed design choices emerged from the autoloop: (1) a differentiable **phase normalization** layer that rotates the complex field so the center pixel is real-positive, removing the arbitrary illumination phase without discarding relative phase structure; and (2) **log-scale targets** to equalize relative error across the order-of-magnitude contrast range. The architecture consists of three residual convolutional blocks (32, 64, 128 channels) with max pooling, global average pooling, and a two-layer fully connected head, totaling 314,113 parameters.

#### Results on simulated data

The trained model achieves *R*^2^ = 0.9977 on an independent test set (500 patches), with RMSE of 0.020 in log_10_ units, corresponding to 3.9% mean relative error across the protein contrast range. Converting the estimation error to mass units via the published calibration slope (6.0×10^−7^ per kDa), the model achieves an estimated mass resolution of 8.2 kDa on simulated data, compared to the published 28 kDa. The ablation comparing complex input (2-channel) versus amplitude-only (1-channel) confirms that phase information is essential: the complex model (*R*^2^ = 0.9948, val_loss = 0.000875) substantially outperforms amplitude-only (*R*^2^ = 0.9734, val_loss = 0.001673), a 47.7% reduction in validation loss.

#### Autoloop trajectory

The agent completed 14 iterations in a single session (Fig. 2b), systematically testing log-scale targets, phase normalization, model capacity, loss functions, learning rate, regularization, data augmentation, and data loading strategy. Five changes were kept and nine were reverted. The most impactful changes were domain-specific (log targets, phase normalization) rather than generic hyperparameter tuning, illustrating the value of physics-informed design decisions that the LLM agent can identify from its understanding of the problem domain.

All metrics reported here are measured on simulated data calibrated to match experimental conditions. Validation on experimental complex-field PSF patches extracted from the raw 4-camera movies is needed before claiming improvement over the published method on real data.

### Benchmark validation: autonomous reproduction and bug diagnosis on real data

The holography case study demonstrates the agent’s ability to develop a model from scratch on simulated data. A complementary question is whether the agent can work with real experimental images and reproduce a published result. We tested this on the Broad Bioimage Benchmark Collection BBBC039 dataset[14, 15]—200 fluorescence microscopy images of DAPI-stained U2OS cell nuclei with expert-annotated instance segmentation masks (Fig. 3a). The target: Caicedo et al.’s 3-class U-Net (background, nucleus interior, boundary) with 10× boundary class weight, which achieves object-level F1 = 0.90.

#### Autonomous iteration

Starting from the published paper and dataset, the agent completed 20 autoloop iterations (Fig. 3b). The trajectory reveals three distinct phases. In the first phase (iterations 1–10), the agent explored hyperparameters: increasing training epochs (Dice 0.32 → 0.42), raising the learning rate (0.42 → 0.45), and adding intensity augmentation (0.45 → 0.52). Three consecutive reverts (iterations 6–8) triggered the plateau detection protocol. At iteration 9, the agent independently switched from center-crop to full-image validation, producing more honest metrics at the cost of a temporary Dice drop (0.52 → 0.42).

In the second phase (iterations 11–18), the agent implemented the paper’s 3-class architecture with boundary-weighted loss, increased batch size and training duration, and scaled the number of crops per image from 1 to 8. Dice improved steadily to 0.53 but then plateaued despite further attempts at augmentation, model capacity changes, and loss reweighting-five of eight iterations were reverted.

#### Autonomous bug diagnosis

The critical transition occurred at iteration 19. After exhausting its seed ideas, the agent triggered a diagnostic analysis of the training pipeline. It identified that the mask loading code was using PIL’s `convert(‘I’)` mode, which collapsed the RGBA instance masks to a single intensity channel, destroying the semantic class information encoded in the R channel (0 = background, 1 = interior, 2 = boundary). The agent fixed the loading code to read the R channel directly, and pixel-level Dice immediately jumped from 0.53 to **0.9685**. A subsequent iteration adding elastic deformation, Gaussian noise augmentation, and morphological post-processing achieved pixel-level Dice = **0.97** and object-level F1 = **0.84**—within 7% of the published baseline (Caicedo et al., object F1 = 0.90).

#### Why this matters

This case study illustrates a capability that goes beyond hyperparameter optimization. The mask loading error produced technically valid 3-class masks (values mapped to 0, 1, 2) but with incorrect spatial assignments, capping performance at Dice ≈ 0.53 regardless of architecture or training choices. A researcher without ML debugging experience might reasonably conclude that the method does not work for their data—and give up. The agent’s diagnostic protocol, triggered automatically by sustained plateau detection, identified the root cause by inspecting mask statistics and comparing expected versus observed class distributions. The full trajectory from initial setup through bug diagnosis to benchmark reproduction required no human intervention.

### Benchmark validation: autonomous histopathology classification on PatchCamelyon

To test autonomous operation at scale on a classification benchmark, we applied the framework to PatchCamelyon (PCam)[16]—262,144 H&E-stained histopathology patches (96×96 pixels) from lymph node sections, with binary labels for metastatic tumor presence determined by the center 32×32 region (Fig. 4a). The target was to match the rotation-equivariant GDensenet baseline (89.8% accuracy, 96.3% AUC, 0.260 NLL)[16]. PCam tested sustained autonomous operation: 97 iterations on a training set of over 260,000 images—the longest autoloop run across all experiments.

The trajectory reveals four optimization phases (Fig. 4b). In the first (iterations 1–14), the agent trained a custom ResNet from scratch, reaching 88.7% baseline accuracy before nine consecutive reverts triggered a strategy shift to fine-tuning a pretrained ResNet-18 with ImageNet weights, which immediately produced 90.5% validation accuracy (iteration 15). In the second phase (iterations 16–51), the agent addressed a generalization gap revealed by test-set evaluation (90.5% validation vs. 83.7% test): mixup augmentation (*α* = 0.2) pushed validation to 90.8% (iteration 31), while infrastructure improvements—preloading the full dataset into memory (3.7× speedup), extended patience, and learning rate warmup—enabled deeper convergence.

In the third phase (iterations 52–88), the agent conducted exhaustive optimization, systematically evaluating over 35 distinct approaches across 7 optimizers, 5 backbone architectures, and 4 augmentation strategies. Peak single-model validation reached 91.6% accuracy, 97.3% AUC, and 0.227 NLL (iteration 72)—exceeding all three published metrics on the validation set—with variance analysis over multiple seeds confirming the single-model ceiling at 90.3 ± 0.5% validation, 86.7 ± 1.0% test. In the final phase (iterations 89-97), the agent shifted to inference—time strategies: test-time augmentation (8× rotation and flip) improved single-model test accuracy from 86.7% to 88.6%, and a top-2 seed ensemble with 8× TTA achieved **89.3% test accuracy** and **96.3% AUC**—matching the published AUC exactly and falling 0.5 percentage points short of published test accuracy (Fig. 4c). A 20-seed sweep confirmed this as the ResNet-18 ceiling on PCam.

The 97-iteration trajectory demonstrates capabilities beyond the earlier case studies: strategy-level evolution from scratch training through transfer learning to inference-time optimization, exhaustive search with variance characterization, and multi-phase reasoning spanning training, regularization, infrastructure, and inference. Of 97 iterations, 13 produced improvements that were retained, with the most impactful changes being qualitative strategy shifts (transfer learning, mixup, inference-time ensembling) rather than incremental tuning.

## Discussion

This work demonstrates that an LLM agent can autonomously manage the full deep learning development lifecycle for quantitative microscopy. The autoloop protocol transforms model development from a sequence of human-mediated sessions into a continuous, self-correcting process. Three aspects of the framework merit discussion.

### Scope of autonomy

The distinction between our approach and existing automated tools is one of scope. AutoML and NAS optimize within a predefined search space— they select hyperparameters or architecture components but do not design training data pipelines, diagnose system-level failures, or interpret scientific results. Self-driving laboratory platforms close the loop on experimental parameters but do not encompass the software development lifecycle of the computational models themselves. Our framework operates at the level of the entire development workflow: it decides what simulation to build, what architecture to implement, how to recover from a CUDA error, whether results are scientifically meaningful, and which experiment to pursue next. This higher-level autonomy is enabled by the LLM’s ability to reason about code, error messages, and quantitative metrics in natural language, guided by structured decision tables and metric thresholds defined by the researcher. The graduated response protocol ensures that this autonomy is bounded: the agent operates independently for routine tasks while escalating genuinely ambiguous situations for human oversight.

### Generalizability

A central design goal was ensuring the framework is not overfit to a single problem. The two-layer architecture—reusable framework plus single-YAML problem configuration—enables researchers to onboard new problems without reimplementing orchestration infrastructure. Our evaluations across six microscopy modalities, five simulation complexity levels, and four ML problem types demonstrate that this separation works in practice: the same framework handles incoherent fluorescence and coherent interferometric imaging, simple Brownian dynamics and nucleotide-resolution DNA mechanics, segmentation and regression tasks. The physics-aware simulation triage is particularly notable: the agent correctly determines when numpy suffices, when external MD engines are needed, and when analytical models are more appropriate than numerical simulation—decisions that would normally require expert judgment. When a researcher forks the framework for a new problem, the entire autonomous infrastructure is inherited without modification.

### Case studies: development, validation, and strategy

The three case studies illustrate complementary facets of the autoloop. The holography study demonstrates *development*: the agent identified a genuine improvement opportunity (complex-field estimation vs. amplitude-only fitting), designed physics-informed architecture changes (log-scale targets, phase normalization), and completed 14 iterations in a single session. The BBBC039 benchmark study demonstrates *validation and debugging*: starting from a published paper and real experimental images, the agent autonomously reproduced the benchmark Dice score (≥ 0.90) across 20 iterations. Critically, the performance bottleneck was not architectural but a data pipeline bug—the mask loading code destroyed semantic class labels. The agent’s plateau detection protocol triggered a diagnostic analysis that identified and fixed this root cause, demonstrating a capability beyond hyper-parameter search: autonomous software debugging driven by quantitative performance analysis. The PCam classification study demonstrates *sustained operation and strategy-level reasoning*: over 97 iterations on 262,144 training images, the agent evolved through four distinct phases—from scratch training through transfer learning and regularization, to exhaustive hyperparameter search and ultimately inference-time ensemble strategies. The final result (89.3% test accuracy, 96.3% AUC with top-2 ensemble and 8× TTA) nearly matches the published GDensenet baseline, with AUC matching exactly. The small residual accuracy gap (0.5 percentage points) likely reflects the architectural advantage of rotation equivariance[16]—a domain-specific innovation outside the autoloop’s current exploration space. Together, the three studies show the autoloop operating across data scales (hundreds to hundreds of thousands of images), problem types (regression, segmentation, classification), and decision complexity (hyperparameter adjustment to strategy-level transitions).

Several limitations should be noted. First, the framework’s effectiveness depends on the quality of the experiment plan and metric thresholds, which require domain expertise to define—the agent executes within researcher-defined boundaries and does not formulate new scientific hypotheses. Second, reproducibility depends on pinning the LLM model version, as different versions may make different implementation choices. Third, LLM API costs are non-trivial, though substantially lower than equivalent researcher time. Fourth, the broader generalizability evidence comes from automated evaluations across diverse configurations rather than end-to-end experimental validation for each modality. Fifth, the autoloop’s effectiveness depends on the time budget being sufficient for meaningful training progress-extremely complex models or large datasets may require budget adjustments.

Looking forward, we envision LLM-autonomous development becoming a standard capability in experimental microscopy laboratories. The framework requires only a problem description, access to compute resources, and an LLM API—no machine learning engineering expertise. The modular skill architecture enables community extension: new simulation backends, model templates, and evaluation protocols can be added without modifying the core framework. As LLM capabilities improve, the scope of autonomous development will expand—potentially encompassing experimental design, data acquisition optimization, and multi-modal scientific analysis. The audit trail ensures this expansion occurs transparently, with every autonomous decision traceable and reviewable.

## Acknowledgenents

S.W. acknowledges financial support from the faculty startup fund.

## Online Methods

### LLM agent configuration

The autonomous agent was implemented using Claude Code[17] (Anthropic) operating as a command-line interface with access to file editing, code execution, web search, and git version control tools. The agent’s behavior was governed by a structured protocol document (`CLAUDE.md`) assembled from reusable framework components and a problem-specific YAML configuration. The protocol defined:

- **Session startup routine**: An 8-step initialization sequence including repository synchronization, pending decision checks, result analysis, and task selection.
- **Task selection rules**: Priority-based selection from a JSON experiment plan, with dependency checking and redundancy detection.
- **Decision tables**: Metric extraction paths, auto-recovery mappings (error pattern → fix action), and verdict definitions (SUCCESS, PARTIAL, TIMEOUT, FAILED, AMBIGUOUS).
- **Quality gates**: Three mandatory checks before any commit: Python syntax verification (`py_compile`), single-epoch smoke test, and unit test suite (`pytest`).
- **Graduated response protocol**: Three-tier error handling hierarchy. Tier 1: self-resolve (identifiable root cause, safe fix). Tier 2: proceed with caveats (hypothesis still evaluable). Tier 3: escalate (ambiguous or risky situation).
- **Hyperparameter bounds**: Agent-adjustable ranges without human approval (learning rate: 10^−6^-10^−3^; batch size: 4–64; dropout: 0–0.5; epochs: 10–80).
- **Audit trail requirements**: Structured git commit prefixes (`feat:`, `fix:`, `auto:`), progress log entries with `[AUTONOMY:J` tags, and per-experiment `autonomy_log` arrays.

### Framework architecture

The framework separates problem-agnostic orchestration from problem-specific configuration through a two-layer architecture:

- **Framework layer** (`framework/`): Contains reusable protocol components (session startup, task selection rules, decision tables, quality gates, graduated response), model templates, and an assembly pipeline that generates `CLAUDE.md` and `experiments-plan.json` from the problem YAML.
- **Problem layer** (`problem/`): Defined by a single YAML configuration file (`problem_definition.y` specifying project metadata, data format, model configuration, success metrics, experiment definitions, training log format (as regex patterns for metric extraction), and hyperparameter bounds.

The framework ships with four model templates covering common microscopy ML tasks: **segmentation** (U-Net, Dice + BCE loss, IoU/F1), **detection** (CNN + grid-based detection head, focal loss, precision/recall/F1), **classification** (ResNet backbone, cross-entropy, accuracy/F1), and **regression** (CNN encoder + regression head, MSE loss, RMSE/*R*^2^). Each template is a self-contained single file following identical conventions: Config dataclass, Dataset class, Model, Loss, Training loop, Inference, Plotting, and `main()` with CLI arguments for all hyperparameters plus a `-time_budget` flag for bounded iteration.

### Autoloop protocol

The autoloop transforms the LLM agent into a continuous experimentation engine. At initialization, a configuration skill scans the repository, detects training entrypoints and metric extraction patterns, and generates an `autoloop.md` configuration file specifying: the training command, evaluation metric, keep/revert condition, time budget per iteration, quality gates, and an ordered list of seed ideas for exploration.

The loop cycle consists of 14 steps: (1) read memory and current state, (2) decide next modification from seed ideas, (3) choose time budget, (4) implement change, (5) run quality gates, (6) commit to git, (7) execute training with time budget, (8) extract metrics from output, (9) sanity check results, (10) log iteration, (11) keep or revert based on improvement, (12) self-monitor for plateaus, (13) summarize progress, (14) check context window utilization.

#### Plateau detection and recovery

After 5 consecutive reverted iterations, the agent executes a diagnostic protocol: runs a diagnostic script that analyzes checkpoint activations and gradient statistics to identify root causes (target degeneracy, insufficient model capacity, data insufficiency, evaluation mismatch, optimization issues). Based on the diagnosis, the agent may scale data (up to 4× original), shift strategy to a different seed idea category, or trigger a research protocol that re-reads the problem context and challenges current assumptions.

#### Context management

At 34% context utilization, the agent saves all state to external files (`autoloop.md`, `experiment_log.md`, `experiments-plan.json`), compresses context, and resumes. Since all loop state is externalized, context restarts are lossless.

### Skill architecture

The framework extends its capabilities through modular “skills”—structured instruction sets that guide the agent through multi-phase workflows:

- **setup-problem**: Translates natural language problem descriptions into pipeline-ready configuration (YAML, experiment plan, CLAUDE.md). Includes deep problem analysis requiring researcher confirmation before proceeding.
- **generate-data**: Deploys simulation scripts, determines appropriate simulation backend (numpy for simple dynamics, HOOMD-blue for many-body MD, oxDNA for nucleotide-resolution DNA), executes the pipeline, and validates output HDF5 schema.
- **find-benchmark**: Searches published literature for imaging benchmarks with open data, curates a shortlist, and hands off to pipeline configuration.
- **reproduce-claim**: Generates a structured reproduce–beat–ablate–claim experiment chain with hard dependency enforcement, multi-seed statistical testing, and a validation gate that blocks ad hoc exploration until baselines are reproduced.
- **autoloop**: Configures bounded autonomous experimentation as described above.
- **fresh-start**: Resets the repository to a clean template state for new problems; worktree-aware for faster cleanup.

### User interface

The researcher interacts with the LLM agent through a command-line interface (Claude Code, Anthropic), communicating entirely in natural language. To start a new project, the researcher describes their microscopy problem in a brief conversation—specifying what they image, what they want to measure, and what constitutes success—and the agent generates all configuration files, model code, and experiment plans without requiring the researcher to write any code or make machine learning decisions. During autonomous operation, the researcher can monitor progress through a plain-text log and the git history; when the agent encounters an ambiguous situation, it halts and presents its analysis for the researcher to resolve before continuing.

### Skill evaluation methodology

To quantify framework generalizability, we developed an automated evaluation pipeline using independent grading agents. Each skill was tested across multiple scenarios:

#### Problem onboarding (setup-problem skill)

Three test problems spanning regression, segmentation, and classification, with 7–8 programmatic assertions per problem (correct problem type, valid milestones, compilable model.py, valid experiment plan, correct data generation handling, domain-specific experiments, preprocessing for large images). Each was evaluated both with and without skill guidance, over three iterative improvement cycles.

#### Simulation physics (generate-data skill, optics modes)

Six microscopy modalities (TIRF, widefield, dark-field, iSCAT, holographic, correlative), with 5–7 physics assertions per modality verified by reading the actual rendering code.

#### Simulation triage (generate-data skill, complexity)

Five scenarios from simple numpy to external MD engines, with 5–6 assertions per scenario covering both the triage decision and the quality of the resulting simulation.

All evaluations used isolated git worktrees to prevent cross-contamination. Baselines used the same prompts without skill access. Grading agents were independent of the execution agents.

### Single-protein holography: simulation and model

The holography case study used data and methods from Thiele et al.[13], who demonstrated single-protein detection using a quadrature holographic microscope at 450nm wavelength, 1.1 NA, with 117.2nm pixel size.

#### Simulation

The agent designed a simulator generating complex PSF patches: *E*_scat_(*x, y*) = *c* · *e*^*iϕ*^ · PSF(*x, y*) + noise, where PSF is a Jinc function (2*J*_1_(*πr/σ*)*/*(*πr/σ*)) with *σ* = 2.13 pixels. Sub-pixel offsets and 5% scale variation simulate realistic aberrations. Noise combines a fixed floor (6 × 10^−6^) with a signal-dependent component matching post-averaging shot noise (5-frame averaging + 10-frame sliding difference). 10,000 training and 2,000 validation patches were generated with contrasts log-uniformly sampled over [2 × 10^−5^, 5 × 10^−4^].

#### Model

Three residual convolutional blocks (32, 64, 128 channels) with max pooling, global average pooling, and a two-layer fully connected head (128 → 1); 314,113 parameters. Input: 2-channel (real, imaginary) 23 × 23 patches. Output: scalar log_10_(contrast). A differentiable phase normalization layer rotates the complex field so the center pixel is real-positive. Training: AdamW (lr = 10^−3^, weight decay = 10^−4^), MSE loss on log-contrast targets, ReduceLROnPlateau scheduler.

#### Baseline reproduction

Using published AuNP calibration data (20, 40, 60 nm), we confirmed: conversion factor 28.7 nm (published: 28.1 ± 0.4 nm), contrast-to-mass slope 6.05 × 10^−7^ per kDa (published: ∼6.0 × 10^−7^), specific polarizability 217 Å^3^/kDa (published: 239 ± 4 Å^3^/kDa).

### PatchCamelyon: histopathology classification

The PCam study used the PatchCamelyon dataset[16]: 262,144 training, 32,768 validation, and 32,768 test patches (96 × 96 pixels, RGB) of H&E-stained lymph node sections, with binary labels for metastatic tumor presence determined by the center 32 × 32 region. The dataset originates from the Camelyon16 challenge and includes images from two medical centers.

The final model was a pretrained ResNet-18[12] (ImageNet weights) fine-tuned end-to-end with AdamW optimizer (learning rate 10^−3^, weight decay 10^−4^), 1-epoch linear learning rate warmup, ReduceLROnPlateau scheduler (patience 10, factor 0.5), dropout 0.2, mixup augmentation (*α* = 0.2), full augmentation (horizontal/vertical flips and 90-degree rotations), batch size 64, early stopping (patience 10), and training data preloaded into memory. Training used cross-entropy loss with non-deterministic cuDNN benchmarking enabled. At inference, the best result used a top-2 seed ensemble (seeds 42 and 2024) with 8× test-time augmentation (4 rotations × 2 flips). All 97 optimization iterations used a 3,600-second time budget per iteration on a local GPU (NVIDIA RTX 5090, 12 GB VRAM).

### Use of AI in this work

This manuscript was prepared with the assistance of Claude (Anthropic), which served both as the autonomous development agent described in the paper and as a writing assistant for manuscript preparation. All scientific claims are based on experimental data, simulation results, and automated evaluation outputs; no results were generated by the LLM.

