## Supplementary material for "Deep learning for microscopy without machine learning expertise": SI

LLM-autonomous development of deep learning models for quantitative microscopy

### Contents

**S1 Problem onboarding evaluation (setup-problem skill) 2**

**S2 Simulation physics evaluation across microscopy modalities 2**

**S3 Simulation complexity triage evaluation 4**

**S4 Combined evaluation summary 5**

**S5 Framework generalizability across problem types 5**

**S6 Holography autoloop iteration details 6**

**S7 Agent-generated code examples 6**

**S8 Evaluation timing and token usage 7**

### S1 Problem onboarding evaluation (setup-problem skill)

Table S1: Problem onboarding evaluation results across three iterative improvement cycles. Each test problem was evaluated both with and without structured skill guidance. Assertions tested: correct problem type, valid milestone thresholds, compilable model.py matching correct template, valid experiment plan schema ($\geq$ 4 experiments), CLAUDE.md with no unfilled placeholders, correct data generation handling, and domain-specific experiments.

| | | With Skill | | Without Skill | |
|---|---|---|---|---|---|
| Iteration | Test Problem | Passed | Total | Passed | Total |
| 1 | AuNP regression | 7 | 7 | 7 | 7 |
| | Nuclear segmentation | 7 | 7 | 6 | 7 |
| | QDot classification | 7 | 8 | 4 | 8 |
| | *Average pass rate* | *95.8%* | | *78.6%* | |
| 2 | AuNP regression | 7 | 7 | 7 | 7 |
| | Nuclear segmentation | 7 | 7 | 7 | 7 |
| | QDot classification | 6 | 8 | 8 | 8 |
| | *Average pass rate* | *91.7%* | | *100%* | |
| 3 (Final) | AuNP regression | 7 | 7 | 6 | 7 |
| | Nuclear segmentation | 8 | 8 | 6 | 8 |
| | QDot classification | 8 | 8 | 8 | 8 |
| | *Average pass rate* | **100%** | | *87.0%* | |

Failure modes in unguided operation included: copying the wrong model template (segmentation template for a classification problem), leaving data generation populated for real-data problems, and missing preprocessing for large images ($1024 \times 1024$). The skill prevented these through explicit decision rules: a template mapping table with verification, a data generation decision tree (real data $\rightarrow$ leave empty), and large-image preprocessing guidance.

### S2 Simulation physics evaluation across microscopy modalities

No modality-specific rendering code was pre-built into the framework. The agent adapted a common baseline (Brownian dynamics + Gaussian PSF) to each modality by rewriting the `render_image()` function based on the researcher's physics description. The specific

Table S2: Physics assertion results for agent-generated simulations across six microscopy modalities. Each assertion was verified by an independent grading agent reading the actual rendering code.

| Modality | Assertions | Passed | Key Physics Verified |
|---|---|---|---|
| TIRF fluorescence | 5 | 5/5 | $\exp(-z/d)$, $d = 150$ nm; PSF $\sigma = 108$ nm |
| Widefield fluorescence | 5 | 5/5 | Born–Wolf defocus; energy-conserving broadening |
| Dark-field scattering | 5 | 5/5 | Mie cross-sections; $d^6 \to d^2$ crossover |
| iSCAT | 6 | 6/6 | $\lvert E_{\mathrm{ref}} + E_{\mathrm{scat}} \rvert^2$; negative contrast |
| Holographic | 6 | 6/6 | $\exp(ikr)/r$; Newton's rings; $z$-encoded fringes |
| Correlative | 7 | 7/7 | 2-channel; 63% conjugated / 37% free |
| **Total** | **34** | **34/34** | |

physics implemented for each modality:

- **TIRF fluorescence**: evanescent wave excitation with exponential $z$-decay ($d \approx$ 150 nm penetration depth) and diffraction-limited PSF ($\sigma = 108$ nm from 532 nm/1.49 NA).
- **Widefield fluorescence**: Born–Wolf defocus model with energy-conserving PSF broadening, $\sigma(z) = \sigma_0\sqrt{1 + (\Delta z/z_R)^2}$.
- **Dark-field scattering**: Mie theory cross-sections with size-dependent intensity scaling (Rayleigh $d^6$ to geometric $d^2$ crossover).
- **Interferometric scattering (iSCAT)**: coherent field addition $I = \lvert E_{\mathrm{ref}} + E_{\mathrm{scat}} \rvert^2$ with $\pi$ phase offset producing negative contrast ($-16\%$ to $-21\%$), mass-dependent polarizability, and spatially correlated speckle noise.
- **Digital holographic microscopy**: spherical wave propagation $\exp(ikr)/r$ with concentric Newton's rings, $z$-encoded fringe spacing, and off-axis tilted reference beam.
- **Correlative (TIRF + dark-field)**: dual rendering pipeline with two particle populations (63% conjugated with tethered dynamics, 37% free Brownian), two-channel output, and co-localization logic.

#### S3 Simulation complexity triage evaluation

Table S3: Simulation complexity triage results. For each scenario, the agent determined the appropriate simulation tool and implemented a working simulation.

| Scenario | Expected Tool | Assertions | Passed |
|---|---|---|---|
| DNA–AuNP assembly | HOOMD-blue | 6 | 6/6 |
| DNA origami folding | oxDNA | 5 | 5/5 |
| Protein diffusion | numpy | 5 | 5/5 |
| Active Janus particles | numpy (RPY) | 5 | 5/5 |
| Vesicle fluctuations | numpy (SH) | 5 | 5/5 |
| **Total** | | **26** | **26/26** |

The five triage scenarios in detail:

1. **DNA–AuNP self-assembly**: the agent correctly identified five physics requirements (Morse potentials, simultaneous pairwise forces, detailed balance, Langevin dynamics, excluded volume) that exceed numpy's capabilities, recommended HOOMD-blue, and implemented a complete simulation script with Morse + WCA potentials and a numpy fallback with documented limitations.

2. **DNA origami folding**: the agent treated oxDNA as a black-box external simulator, generating native input files, invoking the binary via subprocess, parsing trajectory output, and rendering synthetic liquid-AFM images.

3. **Protein membrane diffusion**: correctly chose numpy for simple 2D Brownian motion when the researcher specified "nothing fancy needed."

4. **Active Janus particles**: performed quantitative tradeoff analysis (numpy Rotne–Prager–Yamakawa tensor: $30 \times 30$ mobility matrix at $N = 3$–$15$, trivial Cholesky; HOOMD MPCD: $\sim 10^5$ solvent particles, 100–1000× slower), correctly identifying $N > 100$ as the crossover point.

5. **Vesicle shape fluctuations**: identified three candidate approaches and correctly selected spherical harmonic sampling ($\langle |u_{lm}|^2 \rangle = k_B T / [\kappa(l+2)(l-1)(l(l+1)+\bar{\sigma})]$), rejecting atomistic MD (wrong length scale) and triangulated Monte Carlo (only needed for $\kappa < 2\, k_B T$).

The agent's decisions reveal an emergent triage logic: independent particle dynamics → numpy; many-body interactions at small $N$ → numpy with explicit coupling matrices; large $N$ or researcher-specified engine → external MD package with fallback.

#### S4 Combined evaluation summary

Table S4: Aggregate evaluation results across all test suites.

| Evaluation Suite | Tests | Assertions | Passed | Pass Rate |
|---|---|---|---|---|
| Setup-problem (iter 3, with skill) | 3 | 23 | 23 | 100% |
| Setup-problem (iter 3, baseline) | 3 | 23 | 20 | 87.0% |
| Generate-data: optics modes | 6 | 34 | 34 | 100% |
| Generate-data: simulation triage | 5 | 26 | 26 | 100% |
| **Total (with skill)** | **14** | **83** | **83** | **100%** |

The evaluations covered 11 distinct scientific scenarios spanning 6 imaging modalities (TIRF, widefield, dark-field, iSCAT, holographic, correlative), 5 simulation complexity levels (simple Brownian to nucleotide-resolution DNA mechanics), and 4 ML problem types (segmentation, detection, classification, regression).

#### S5 Framework generalizability across problem types

To demonstrate that the framework is not overfit to a single domain, we tested problem onboarding, model template selection, and configuration generation across four ML problem types:

**Segmentation** (nuclear segmentation from DAPI fluorescence): the agent selected the U-Net template with Dice + BCE loss, configured IoU and F1 metrics with thresholds, designed experiments addressing the key challenge of separating touching nuclei, and correctly handled $1024 \times 1024$ input images by configuring patch-based preprocessing (resize to 128, batch size 8).

**Detection** (nanoparticle detection in dark-field images): the agent selected the CNN detection template with focal loss, configured precision/recall/F1 metrics, and designed an experiment progression from baseline detection through sub-pixel localization.

**Classification** (quantum dot counting from widefield fluorescence): the agent selected the ResNet classification template, correctly left the `data_generation` section empty (real data provided as 10K HDF5 crops), and designed experiments addressing class imbalance across five count categories.

**Regression** (AuNP hydrodynamic sizing from scattering microscopy): the agent selected the regression template with MSE loss, configured RMSE and $R^2$ metrics, and correctly populated the simulation section for Langevin dynamics with Mie theory rendering.

Each template is a self-contained single Python file containing configuration, dataset, model, loss, training loop, inference, and `main()` with CLI arguments—ensuring that the agent can modify any component without dependency conflicts. Failures in unguided mode included copying the wrong model template (segmentation for classification) and leaving data generation populated for real-data problems—errors prevented by the skill's explicit decision rules.

### S6 Holography autoloop iteration details

The agent completed 14 iterations in a single session for the holography case study. Table S5 summarizes each iteration.

The most impactful changes were domain-specific (log-scale targets, phase normalization) rather than generic hyperparameter tuning. The ablation in iteration 13 confirmed that complex-valued input provides 47.7% lower validation loss than amplitude-only input.

### S7 Agent-generated code examples

Representative code snippets illustrating: (a) the holography PSF simulator designed by the agent, including Jinc function generation and noise model; (b) the phase normalization layer that removes nuisance phase while preserving relative structure; (c) an autoloop keep/revert decision based on metric comparison. Each example includes the agent's

Table S5: Autoloop iteration log for the holography case study. Changes that improved the best metric were kept; others were reverted via git.

| Iteration | Change | Outcome | $R^2$ |
|---|---|---|---|
| 1 | Baseline (linear contrast targets) | Keep | 0.94 |
| 2 | Log-scale targets | Keep | 0.97 |
| 3 | Phase normalization layer | Keep | 0.98 |
| 4 | Increase model capacity (128 ch) | Revert | – |
| 5 | Huber loss | Revert | – |
| 6 | Higher learning rate ($10^{-3}$) | Keep | 0.99 |
| 7 | Dropout (0.2) | Revert | – |
| 8 | Weight decay ($10^{-3}$) | Revert | – |
| 9 | Data augmentation (rotation) | Revert | – |
| 10 | Cosine annealing scheduler | Revert | – |
| 11 | RAM preloading | Keep | 0.99 |
| 12 | Aggressive LR ($3 \times 10^{-3}$) | Revert | – |
| 13 | Ablation: amplitude only | Revert | 0.97 |
| 14 | Ablation: no phase norm | Revert | – |

reasoning as recorded in the experiment log. Code is available in the accompanying repository.

### S8 Evaluation timing and token usage

Table S6: Token consumption and execution time for skill evaluations and the holography case study.

| Task | Avg. Tokens | Avg. Duration (s) |
|---|---|---|
| Setup-problem (with skill) | 122,000 | 446 |
| Setup-problem (baseline) | 80,000 | 471 |
| Generate-data: optics modes | 62,600 | 348 |
| Generate-data: simulation triage | 60,300 | 392 |
| Holography autoloop (14 iterations) | – | single session |

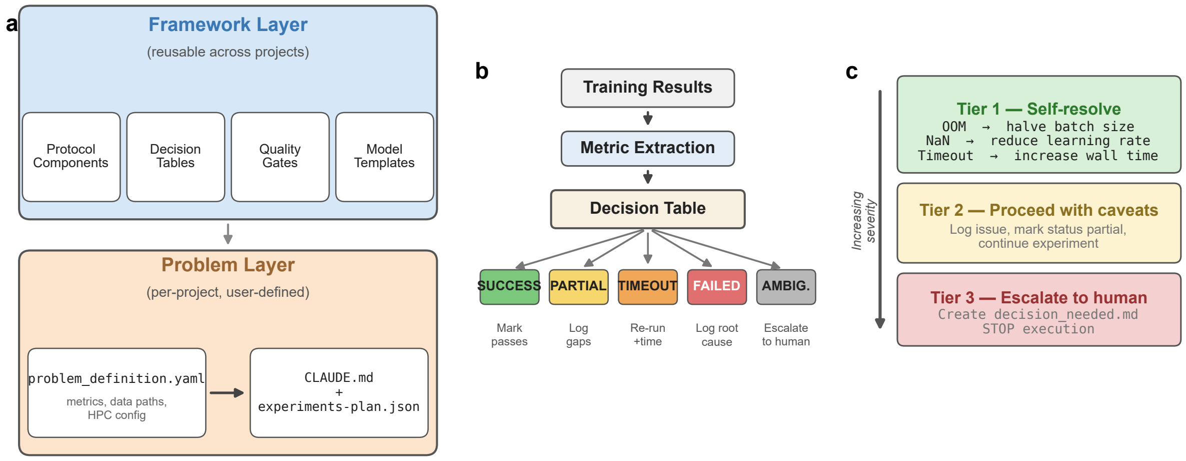


Figure Extended Data 1: **Two-layer framework architecture.** **(a)** The framework separates reusable infrastructure (protocol components, SLURM scripts, model templates) from problem-specific configuration (single YAML file). An assembly pipeline generates the agent protocol and experiment plan. **(b)** Decision tables map training metrics to structured verdicts. **(c)** The graduated response protocol defines three tiers of error handling, ensuring autonomous operation for routine tasks while escalating ambiguous situations for human oversight.

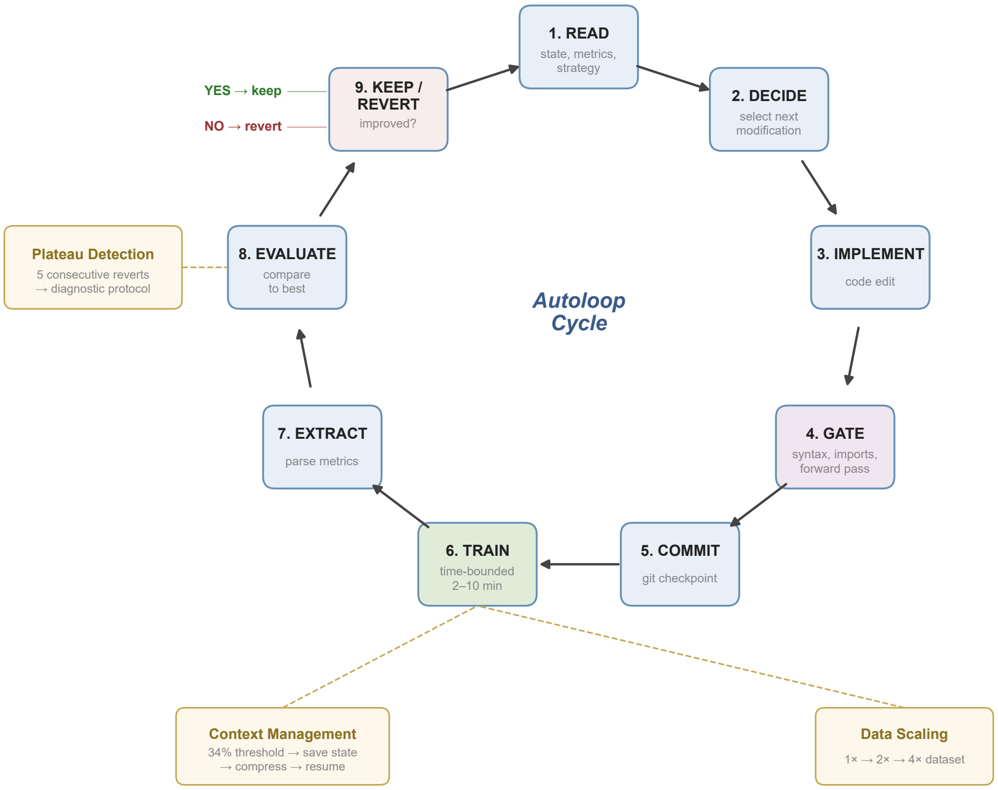


Figure Extended Data 2: **The autoloop cycle in detail.** The agent executes an indefinite cycle of nine steps: read state, decide modification, implement, gate (quality checks), commit (git checkpoint), train (time-bounded), extract metrics, evaluate against best, and keep or revert. Three anti-stalling mechanisms—plateau detection (5 consecutive reverts triggers diagnostic protocol), automatic data scaling (up to 4×), and context window management (34% threshold)—ensure sustained autonomous operation.
